# A tunable immune-evasion function of Zika virus NS5 governs viral fitness and pathogenesis

**DOI:** 10.64898/2025.12.28.696729

**Authors:** Yu Zhang, Yaqing Zhang, Chengyu Liu, Wenlin Ren, Chenhao Li, Yongqi Guo, Leyi Bai, Peigang Wang, Jing An, Jun Liu, Qiang Ding

## Abstract

Type I interferon (IFN) signaling is a central antiviral defense, with STAT2 driving the expression of IFN-stimulated genes (ISGs) to restrict viral infection. Many flaviviruses, including Zika virus (ZIKV), evade this pathway through non-structural protein 5 (NS5)-mediated ubiquitination and proteasomal degradation of STAT2, yet the contribution of this immune evasion strategy to viral fitness and pathogenesis remains incompletely defined. Here, using a multi-step mutational scanning strategy that jointly assessed STAT2 degradation and viral RNA replication, we identified a single NS5 residue, L162, as uniquely permissive to substitution that abolishes STAT2 degradation without compromising intrinsic replication functions. Substitution of L162 with alanine or glycine (L162A or L162G) selectively disrupted NS5 recruitment of the ZSWIM8-CUL3 E3 ubiquitin ligase, thereby preserving STAT2 stability and restoring IFN signaling. Recombinant ZIKV carrying these mutations exhibited reduced replication and enhanced ISG induction in human cells, defects fully rescued by STAT2 knockout. In A129 mice (type I IFN receptor deficient), mutant and WT viruses replicated comparably, but in human STAT2 knock-in mice, ZIKV-NS5^L162G^ exhibited markedly reduced viral loads and disease. Notably, infection with ZIKV-NS5^L162G^ elicited robust neutralizing antibody and T cell responses that conferred protection against WT challenge. Together, these findings establish NS5-mediated STAT2 degradation as a central determinant of ZIKV immune evasion, viral fitness and pathogenesis, and highlight disruption of NS5-STAT2 antagonism as a promising strategy for antiviral intervention and rational attenuation.

## Introduction

Zika virus (ZIKV) is a mosquito-borne flavivirus within the family *Flaviviridae* that has caused millions of infections worldwide and emerged as a major public health threat (1, 2). Other medically important flaviviruses include dengue virus (DENV), yellow fever virus (YFV), Japanese encephalitis virus (JEV), and West Nile virus (WNV), which together account for significant global morbidity and mortality (3). ZIKV infection is associated with severe neurological complications. In fetuses and neonates, it can cause congenital Zika syndrome characterized by microcephaly, ventriculomegaly, and other brain malformations, while in adults it has been linked to Guillain-Barre syndrome (4).

Like other flaviviruses, ZIKV is an enveloped, single-stranded, positive-sense RNA virus with an approximately 11-kb genome that encodes a single polyprotein, which is co- and post-translationally processed into three structural proteins (C, prM and E) and seven nonstructural proteins (NS1-NS5) (5). The capsid (C) packages the viral RNA genome, the precursor membrane protein (prM) and envelope (E) protein mediate virion assembly and receptor binding/fusion, respectively. Among the nonstructural proteins, NS1 plays roles in RNA replication and immune evasion, NS2A and NS2B function in virion assembly and polyprotein processing, NS3 provides protease and helicase activities, NS4A and NS4B remodel host membranes and antagonize innate immune signaling, and NS5 serves as the viral RNA-dependent RNA polymerase and methyltransferase while also acting as a potent antagonist of type I interferon (IFN) signaling (6). Together, these nonstructural proteins orchestrate viral RNA replication, virion production, and modulation of host antiviral responses.

IFN signaling is a critical first line of defense against viral infection. Viral nucleic acids are sensed by distinct pattern-recognition receptors (PRRs) that activate convergent pathways leading to IFN production. Cytosolic RIG-I-like receptors (RIG-I, MDA5) detect viral RNA and signal through the adaptor MAVS (7–10), whereas endosomal Toll-like receptors (TLR3, TLR7/8) signal through TRIF or MyD88 (11, 12). Cytosolic DNA is recognized by the cGAS sensor, which activates the adaptor STING (13–15). These pathways converge on the kinases TBK1 and IKKε, which phosphorylate IRF3 and IRF7 to drive type I IFN transcription (16, 17). The secreted IFNs then bind the type I IFN receptor (IFNAR) on neighboring cells, activating the JAK-STAT pathway and leading to phosphorylation of STAT1 and STAT2. Together with IRF9, they form the ISGF3 complex, which translocates to the nucleus to induce hundreds of IFN-stimulated genes (ISGs) that establish an antiviral state (18).

Flaviviruses have evolved multiple, complementary strategies to evade type I IFN responses, thereby promoting efficient replication and pathogenesis (19). They avoid immune recognition by cloaking their RNA through 2’-O-methylation of the viral cap by NS5, preventing detection by MDA5 (20, 21). NS1 interferes with TLR3 signaling, and subgenomic flavivirus RNA (sfRNA) binds the ubiquitin ligase adaptor TRIM25, blocking its deubiquitylation and thereby suppressing RIG-I-mediated signaling (22, 23). DENV and ZIKV NS3 proteins interact with 14-3-3 chaperones to prevent RIG-I activation (24), DENV NS2B promotes lysosomal degradation of the DNA sensor cGAS to block detection of mitochondrial DNA (25), and the NS2B/NS3 protease cleaves STING to disable cGAS/STING signaling (26, 27). NS4B further inhibits TBK1 activation and JAK/STAT phosphorylation, reducing ISG expression (28). Among these mechanisms, NS5 is a potent antagonist of IFN signaling that engages distinct host ubiquitin ligases across flaviviruses. ZIKV NS5 hijacks the ZSWIM8-CUL3 E3 ligase complex to mediate STAT2 ubiquitination and proteasomal degradation, whereas DENV NS5 recruits UBR4 to achieve the same outcome (29, 30). Notably, ZIKV and DENV NS5 cannot degrade murine STAT2 (31, 32), explaining why immunocompetent mice are largely resistant to infection (33), whereas type I IFN receptor-deficient or human STAT2 knock-in mice are highly susceptible and serve as crucial models to study viral pathogenesis (34, 35). Despite these insights, the precise molecular determinants within ZIKV NS5 that mediate ZSWIM8-CUL3 recruitment and their direct contribution to infection and pathogenicity *in vivo* remain incompletely defined.

Live-attenuated vaccines have proven highly effective for other flaviviruses, such as YFV-17D (36, 37) and JEV-SA14-14-2 (38, 39), which induce robust and durable immunity. Attenuation of YFV-17D is largely attributed to its reduced ability to evade innate immune responses, particularly type I IFN signaling, while retaining sufficient replicative capacity to trigger strong, long-lasting immunity (40, 41). Several ZIKV live-attenuated vaccine candidates have been developed, incorporating mutations in the E protein glycosylation site (42), deletions in C or NS1 (43, 44), or insertions in the 5’UTR (45). However, these genetic modifications directly impair viral replication, leading to poor viral growth in cell culture and consequently low yields, which complicates large-scale vaccine production.

In this study, we employed a multistep functional screening strategy to identify ZIKV NS5 variants that selectively lose the ability to degrade STAT2 while retaining intrinsic viral RNA replication function. This approach revealed residue L162 as uniquely permissive to substitutions that disrupt STAT2 degradation without compromising NS5-dependent genome replication. Recombinant ZIKV carrying L162 substitutions exhibited STAT2-dependent restriction of infection and diminished pathogenicity in immunocompetent hosts, yet remained highly immunogenic, eliciting robust neutralizing antibody and T cell responses that conferred complete protection against subsequent wild-type (WT) ZIKV challenge. Together, our findings demonstrate that NS5-mediated STAT2 degradation is a major determinant of ZIKV immune evasion and pathogenesis, and provide proof of concept for rational design of live-attenuated vaccines by targeting viral immune antagonism.

## Results

### Identification of ZIKV NS5 mutants defective in STAT2 degradation but competent for viral RNA replication

ZIKV NS5 serves as both the viral methyltransferase and RNA-dependent RNA polymerase, directly mediating viral RNA synthesis (46). Beyond its replication role, NS5 antagonizes IFN signaling by hijacking the host ZSWIM8-CUL3 E3 ligase to promote STAT2 ubiquitination and proteasomal degradation, thereby suppressing ISG induction (**Fig. S1A**) (29, 32). To dissect the contribution of NS5-mediated immune evasion to viral infection and pathogenesis, we sought to identify NS5 mutants that lose the ability to degrade STAT2 while retaining viral genome replication functions (**Fig. S1A**).

We first employed an IFN-stimulated response element (ISRE) luciferase reporter assay (47) to screen for NS5 mutants defective in inhibiting IFN signaling (**Fig. S1B**). We individually generated a panel of 151 alanine-scanning NS5 mutants, in which six consecutive residues were replaced with alanines across the 903-amino-acid NS5 protein (**Fig. 1A**). Thirty-three mutants, corresponding to 198 amino acid positions, lost this inhibitory activity (**Fig. 1A** and **Fig. S1C**, red columns). To refine these regions, we performed a secondary screen using overlapping two- to four-residue alanine substitutions within each of the 33 candidate segments. STAT2 degradation was quantitatively assessed using a STAT2-mCherry reporter system (**Table S1**), narrowing the critical determinants to 149 individual residues required for NS5-mediated STAT2 degradation (**Fig. 1A**).

**Figure 1.**
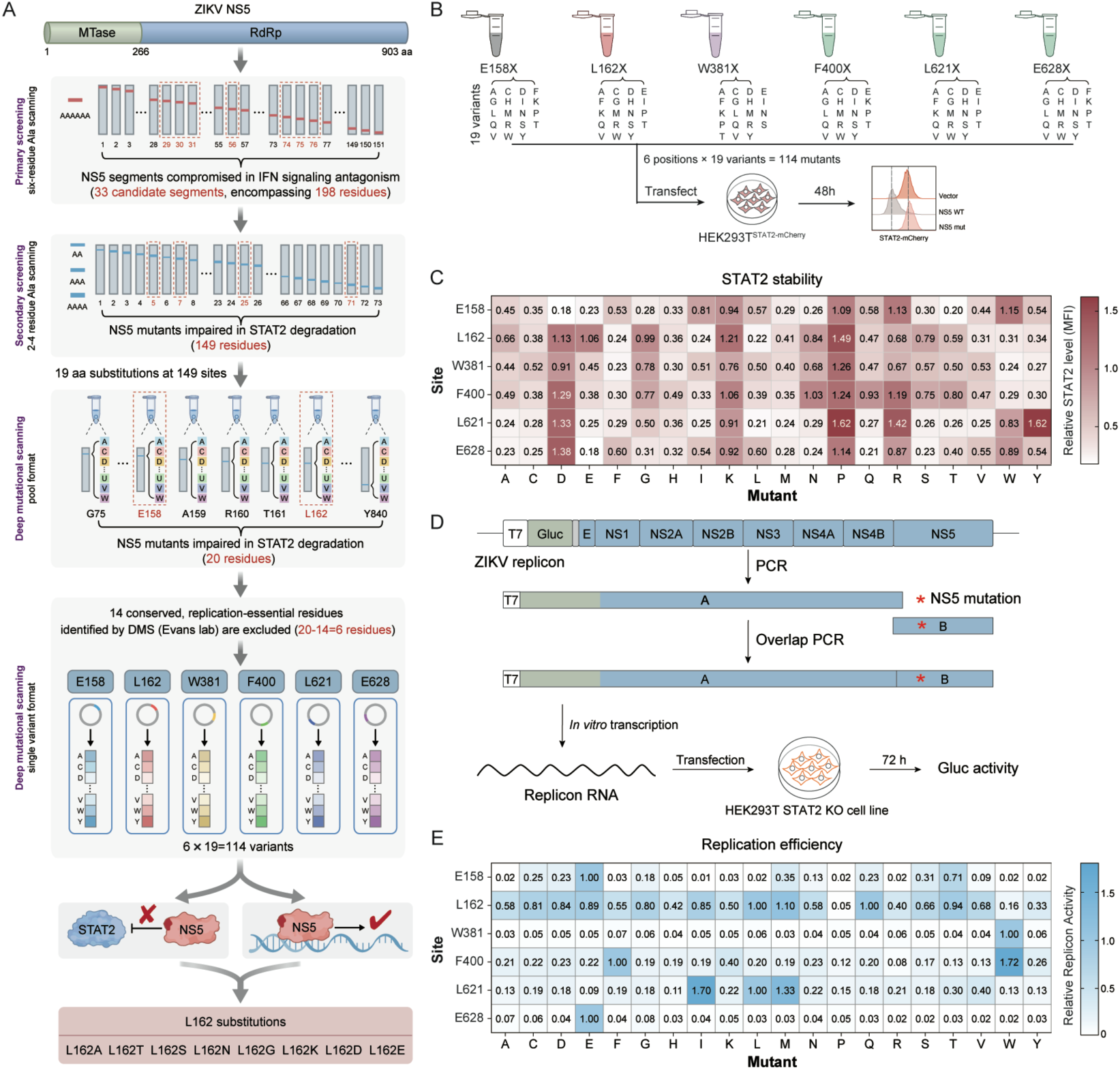
A multistep functional screening strategy identifies NS5 residue(s) required for STAT2 degradation independently of viral RNA replication. **(A)** Schematic overview of the multistep screening strategy used to identify NS5 residues required for STAT2 degradation but not viral RNA replication. An initial alanine-scanning screen across the 903-amino acid NS5 protein was performed using an ISRE luciferase reporter assay. Candidate regions were subsequently refined by secondary alanine scanning combined with STAT2-mCherry-based flow cytometric analysis. Selected residues were then subjected to deep mutational scanning, followed by parallel evaluation of STAT2 degradation and viral RNA replication using a ZIKV replicon system. (B-C) Functional profiling of single-amino acid substitutions at six candidate NS5 residues (E158, L162, W381, F400, L621 and E628; 6 residues × 19 substitutions=114 variants). STAT2-mCherry reporter cells were seeded at 2 × 10^5^ cells per well and individually transfected with 500 ng of plasmid encoding each NS5 variant. STAT2 abundance was quantified by flow cytometry at 48 h post-transfection. Mean fluorescence intensity (MFI) values were normalized to vector control (B). A heat map summarizes STAT2 degradation activity across all 114 variants, with color intensity reflecting relative STAT2-mCherry abundance (C). (D-E) Single-amino acid substitutions were introduced into a Gaussia luciferase (Gluc)-expressing ZIKV replicon by PCR assembly. *In vitro*-transcribed replicon RNA (500 ng) was transfected into HEK293T STAT2 KO cells (2 × 10^5^ cells per well). Replication efficiency was measured by Gluc activity in culture supernatants at 72 h post-transfection (D) and is shown as relative luciferase activity normalized to WT NS5 (E).

We next conducted deep mutational scanning of these 149 residues, substituting each position with all 19 alternative amino acids and screening the resulting variants in pooled format for loss of STAT2 degradation activity (**Fig. S2A-C**). This analysis identified 20 residue positions whose mutation markedly impaired STAT2 degradation (**Fig. S2C,** red columns). Comparison with prior ZIKV genome deep mutational scanning datasets revealed that 14 of these residues (R201, Y301, Y306, H443, C448, C451, E488, L493, F492, W498, L636, D734, Y768 and W815) are highly conserved and indispensable for viral replication (48). Because mutations at these positions impair viral replication, they were excluded from further analysis. The remaining six candidate residues (E158, L162, W381, F400, L621 and E628) were subsequently subjected to detailed mutational characterization. Each residue was individually substituted with all 19 alternative amino acids, yielding a panel of 114 NS5 variants. STAT2 degradation activity was quantified using the STAT2-mCherry reporter cells, revealing a spectrum of partial to complete loss of activity (**Fig. 1B-C**). In parallel, each variant was introduced into a ZIKV replicon and assessed for replication competence in STAT2-knockout HEK293T cells (**Fig. 1D-E**). Notably, only substitutions at residue L162 (including L162A, L162T, L162S, L162N, L162G, L162K, L162D, and L162E) markedly abolished NS5-mediated STAT2 degradation while largely preserving viral RNA replication.

Collectively, this multistep screening strategy identifies residue L162 in NS5 as a unique determinant whose mutations selectively disrupt STAT2 degradation without directly compromising the intrinsic replication function of NS5. These L162-directed variants provide genetic tools to dissect the contribution of NS5-mediated STAT2 degradation to ZIKV infection and pathogenesis.

### NS5 L162 mutations disrupt NS5-ZSWIM8 interaction, restores STAT2 stability, reactivates IFN signaling, and restrict ZIKV infection

After identifying residue L162 as a critical determinant of NS5-mediated STAT2 degradation, we selected two representative substitutions, L162A and L162G, for subsequent study. These mutations were engineered into the ZIKV Dakar-41525 strain genome (GenBank: MG758785.1) (49) and *in vitro*-transcribed viral genomic RNAs were electroporated into Vero cells to rescue two recombinant viruses, ZIKV-NS5^L162A^ and ZIKV-NS5^L162G^, by reverse genetics (**Fig. 2A**). Upon infection of A549 cells, both mutant viruses failed to induce STAT2 degradation compared with WT ZIKV (**Fig. 2B**). Co-immunoprecipitation assays showed that WT and mutant NS5 proteins associated comparably with STAT2, whereas the L162A and L162G substitutions markedly impaired NS5 interaction with ZSWIM8, with the L162G mutant displaying the most pronounced defect (**Fig. 2C**). Consistent with this finding, structural analysis of the NS5-STAT2 complex (50) places residue L162 outside the STAT2 interaction interface (**Fig. 2D**), and its substitution does not affect NS5-STAT2 association. Together, these results indicate that L162 is required for efficient NS5-ZSWIM8 association and that disruption of this interaction abolishes NS5-mediated STAT2 degradation (**Fig. 2C-D**).

**Figure 2.**
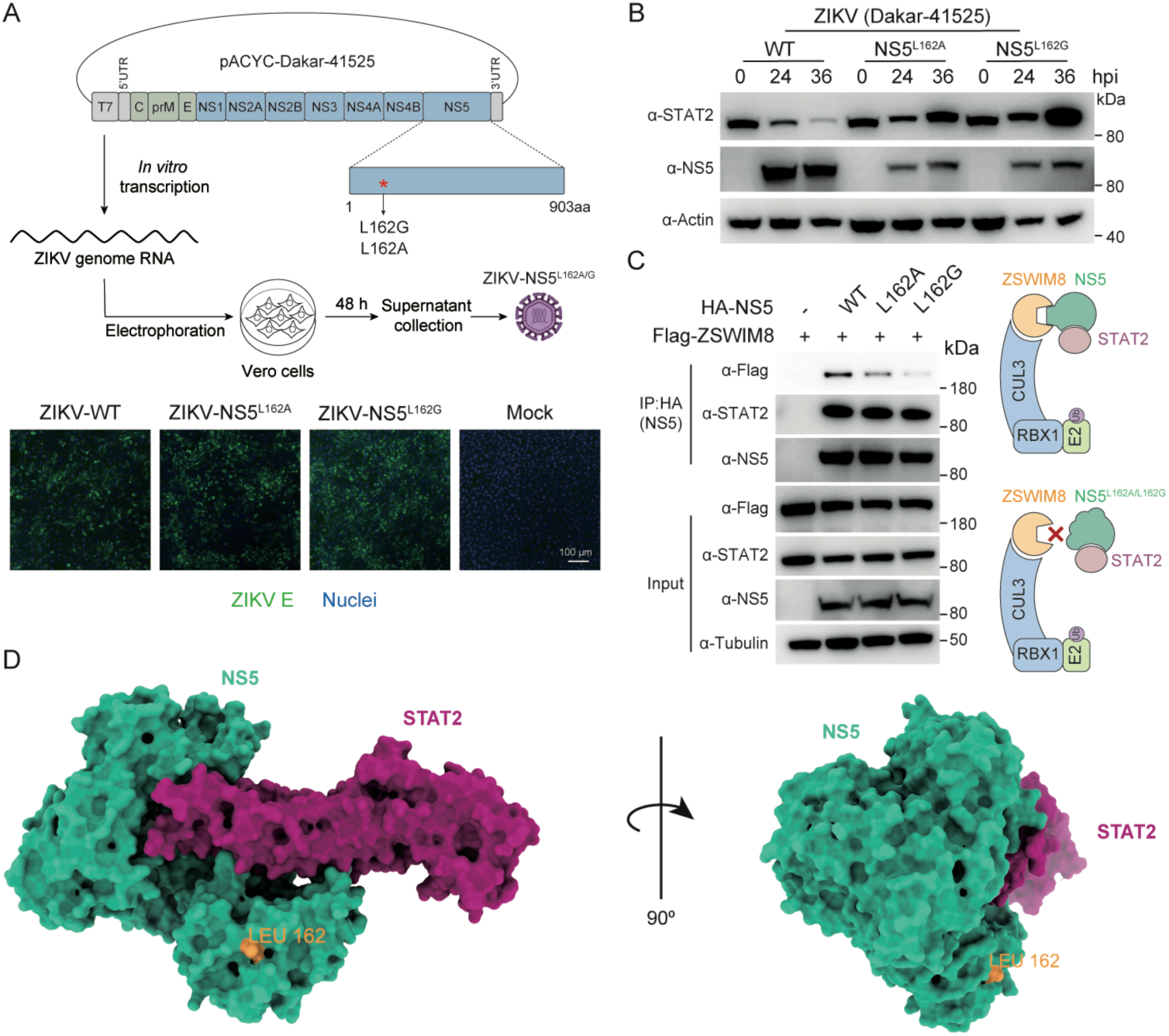
Generation of ZIKV NS5 L162 mutant viruses and analysis of NS5-STAT2 and NS5-ZSWIM8 interactions. (**A**) Schematic of recombinant ZIKV generation by reverse genetics. L162A or L162G substitutions were introduced into the NS5 coding region of the ZIKV Dakar-41525 infectious clone (GenBank: MG758785.1). *In vitro*-transcribed full-length viral genomic RNAs (10 μg) were electroporated into Vero cells (8×10^6^ cells) and culture supernatants were collected 48 h post-electroporation to generate recombinant virus stocks. Immunofluorescence images show ZIKV E protein (green) and nuclei (blue) in Vero cells infected with WT ZIKV, ZIKV-NS5^L162A^, ZIKV-NS5^L162G^, or mock control. Scale bar, 100 μm. (**B**) A549 cells were infected with WT ZIKV or NS5 mutant viruses at an MOI of 0.1 and harvested at the indicated time points post-infection. Cell lysates were analyzed by immunoblotting using antibodies against STAT2 and NS5; β-actin served as a loading control. (**C**) HEK293T cells (1×10^6^ cells) were co-transfected with plasmids expressing HA-tagged WT or mutant NS5 (L162A or L162G; 1.5μg each) together with Flag-tagged ZSWIM8 (1.5μg). At 72 h post transfection, cell lysates were subjected to anti-HA immunoprecipitation, followed by immunoblotting for co-precipitated STAT2 and Flag-ZSWIM8. Input controls are shown. (**D**) Structural mapping of residue L162 in the ZIKV NS5-STAT2 complex (PDB: 6WCZ). L162 is highlighted on the NS5 structure. All experiments were performed at least three times with similar results and representative images are shown.

We next assessed the impact of these mutations on viral infection *in vitro*. WT ZIKV and NS5 mutant viruses were used to infect WT A549 cells (sgNC) or STAT2 knockout cells (sgSTAT2) at an MOI of 0.01 (**Fig. 3A**). In WT A549 cells, WT ZIKV induced pronounced cytopathic effects (CPE), whereas both NS5 mutant viruses produced substantially weaker CPE, with ZIKV-NS5^L162G^ showing the greatest reduction (**Fig. 3B, top**), consistent with its more severe defect in ZSWIM8 binding (**Fig. 2C**). In contrast, all three viruses caused comparable CPE in A549-STAT2 KO cells (**Fig. 3B, bottom**). Viral titration assays performed on Vero cells revealed reduced viral yields in WT A549 cells, with titers of 8.9 × 10^5^ FFU/mL for ZIKV-NS5^L162A^ and 6.0 × 10^5^ FFU/mL for ZIKV-NS5^L162G^, corresponding to an approximately 7- to 13-fold reduction compared to WT ZIKV (7.7 × 10^6^ FFU/mL) (**Fig. 3C; Fig. S3A**). By contrast, all three viruses achieved similar titers in A549-STAT2 KO cells (**Fig. 3C**). These results indicate that the reduced viral infection for NS5 mutant viruses in WT A549 cells are STAT2-dependent and not due to an intrinsic defect in viral RNA replication. Serial passage of WT and NS5 mutant viruses in A549 or Vero cells (up to 10 passages) revealed no changes at the NS5 mutation site, indicating genetic stability of the L162A and L162G mutants **(Fig. S3B-C**).

**Figure 3.**
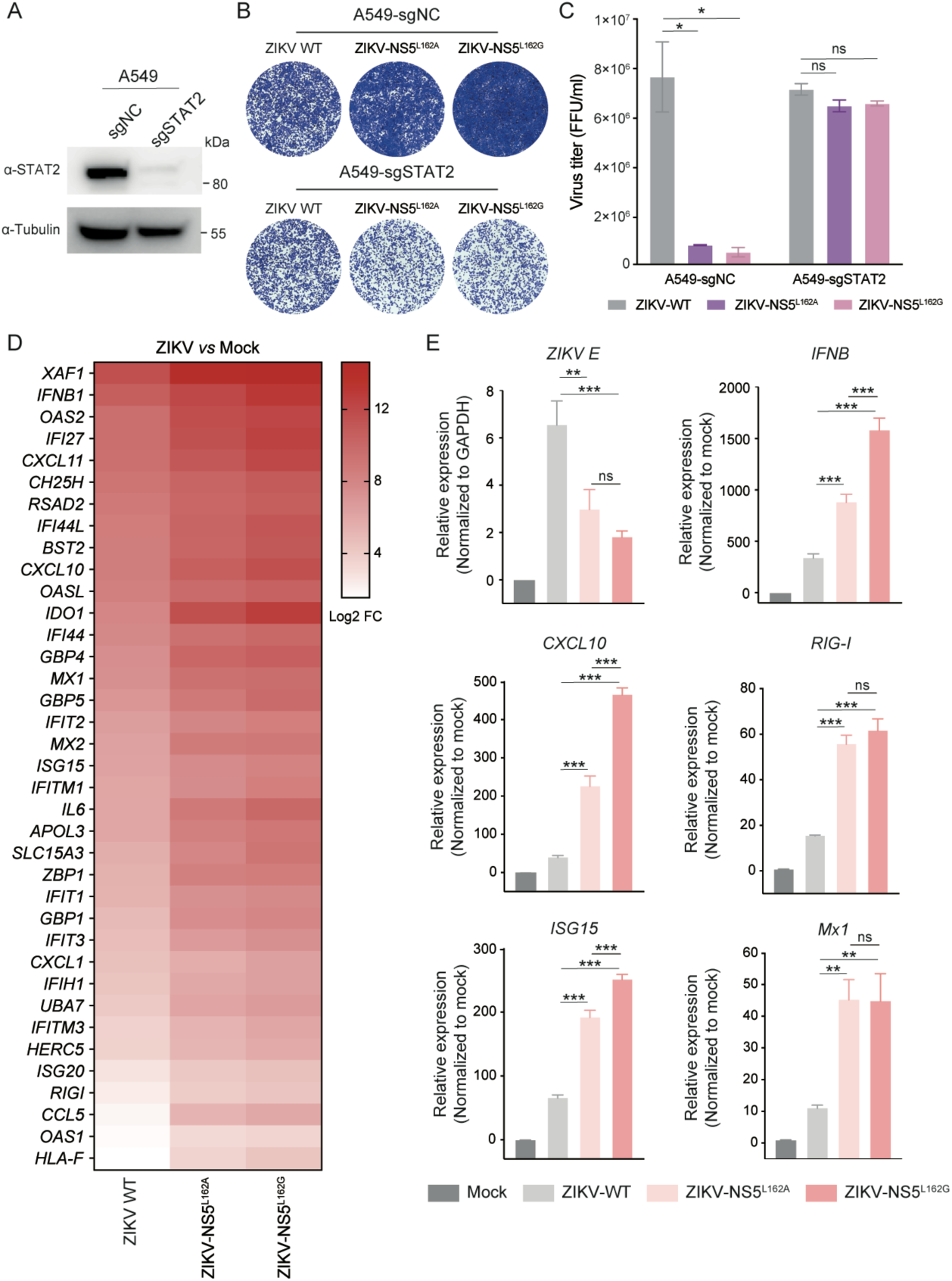
Characterization the infection of ZIKV-NS5^L162A^ and ZIKV-NS5^L162G^ recombinant viruses *in vitro*. (**A**) A549 cells were transduced with lentiviruses encoding Cas9 and either a non-targeting (sgNC) or STAT2-targeting sgRNA (sgSTAT2). Following puromycin selection, STAT2 expression was assessed by immunoblotting. Tubulin served as the loading control. (**B**) A549-sgNC or A549-sgSTAT2 cells were infected with WT ZIKV, ZIKV-NS5^L162A^, or ZIKV-NS5^L162G^ (MOI=0.01). After 2 h, cells were washed with PBS and incubated in fresh medium. At 48 h post-infection (hpi), cell monolayers were fixed and stained with crystal violet to visualize cytopathic effect (CPE) (**B**), while culture supernatants collected in parallel were subjected to focus-forming assay (FFA) on Vero cells to determine viral titers (**C**). Viral titers are presented as focus-forming units per milliliter (FFU/ml). (**D-E**) WT A549 cells were infected with WT ZIKV or NS5 mutant viruses (MOI = 0.05). Total RNA was extracted at 48 hpi and subjected to RNA-seq analysis. Heat map displays |Log₂ fold change| values for upregulated genes associated with immune and antiviral response pathways in virus-infected cells relative to mock-infected controls (**D**). In paralleled, the expression of *IFN-β, CXCL10, RIG-I, ISG15, MX1* and viral *E* gene was quantified by RT-qPCR. Data were normalized to GAPDH and expressed relative to mock-infected controls (**E**). Bars represent mean±SEM. Statistical significance was determined using one-way ANOVA with Tukey’s post hoc test (*p < 0.05; **p < 0.01; ***p < 0.001; ns, not significant).

To further investigate the impact of NS5 mutations on IFN signaling, WT A549 cells were infected with WT or the NS5 mutant viruses (MOI=0.05), and total RNA was collected at 48 hpi. Transcriptome profiling by RNA sequencing revealed a broad and robust upregulation of interferon-stimulated genes (ISGs) in cells infected with the NS5 mutant viruses compared with WT ZIKV, with both the magnitude and breadth of ISG induction being markedly enhanced (**Fig. 3D**). Gene Ontology (GO) enrichment analysis showed that genes upregulated during NS5 mutant virus infection were significantly enriched in antiviral and interferon-related pathways (**Fig. S4**). RT-qPCR analysis further validated the RNA-seq findings, confirming significantly elevated expression of IFN-β and multiple representative ISGs, including *CXCL10, RIG-I, ISG15*, and *MX1*, in mutant virus-infected cells, particularly in response to ZIKV-NS5^L162G^ (**Fig. 3E**). In parallel, viral RNA levels were substantially reduced in cells infected with the NS5 mutant viruses relative to WT ZIKV, consistent with attenuated viral replication under conditions of intact IFN signaling.

Collectively, these findings demonstrate that disruption of NS5-mediated STAT2 degradation compromises ZIKV immune evasion, leading to enhanced IFN signaling, elevated ISG induction, and STAT2-dependent restriction of viral infection *in vitro*.

### ZIKV-NS5^L162G^ virus exhibits STAT2-dependent restriction of infection and pathogenicity in vivo

To investigate the *in vivo* relevance of NS5-mediated STAT2 degradation, we compared the infection and pathogenicity of WT ZIKV and the ZIKV-NS5^L162G^ mutant in two complementary mouse models that differ in their type I IFN competence. We first utilized A129 mice, which lack type I IFN receptors and are widely used as an immunodeficient model for flavivirus infection (34). A129 mice were inoculated subcutaneously with WT ZIKV or ZIKV-NS5^L162G^ (Dakar strain). Body weight was monitored daily, blood was collected at the indicated time points, and viral RNA loads were quantified in organs at day 4 post-infection and at the experimental endpoint (**Fig. 4A**). Both WT ZIKV and NS5^L162G^-infected mice exhibited significant weight loss compared with mock-infected controls (**Fig. 4B**), and 100% mortality was observed within six days in both groups (**Fig. 4C**). Viral RNA levels measured in the brain, heart, spleen, and blood at days 4 and experimental endpoint were comparable between WT and NS5^L162G^-infected mice (**Fig. 4D-E**), confirming similar tissue tropism and viral dissemination under IFN-deficient conditions. Although slightly lower viremia was noted in NS5^L162G^-infected mice at later time points (4-7 dpi) (**Fig. 4E**), this likely reflects biological variability due to the limited sample size. These results demonstrate that the NS5^L162G^ mutation does not impair viral infection or pathogenicity in the absence of type I IFN signaling *in vivo*.

**Figure 4.**
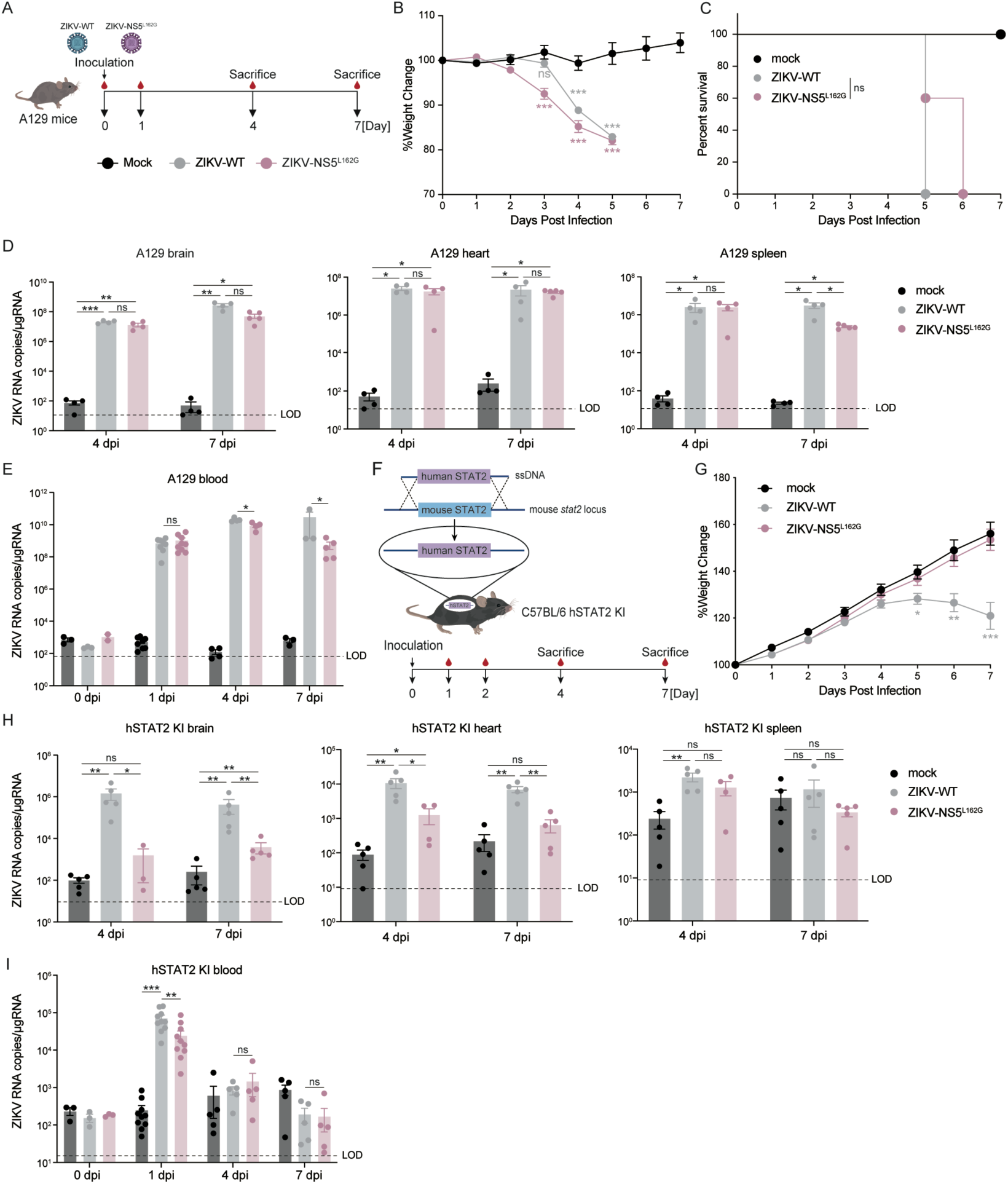
Characterization of ZIKV-NS5^L162G^ mutant virus in IFN-deficient and immunocompetent mouse models. (**A**) Schematic of the experimental design. Six- to eight-week-old A129 mice were inoculated via footpad injection with WT ZIKV or ZIKV-NS5L162G (1× 10^5^ FFU per mouse) or mock treated with PBS (n = 8 for mock, n = 8 for WT ZIKV and n = 9 for ZIKV-NS5^L162G^). Body weight was monitored daily and blood, brain, heart and spleen were collected at 4 and 7 days post-infection (dpi) for viral RNA quantification. (**B-C**) Body weight changes and survival curves of A129 mice following infection with WT ZIKV or ZIKV-NS5^L162G^ virus, or mock infection. (**D-E**) Blood and indicated tissues from infected mice were harvested at 4 dpi and experimental endpoint (5-7 dpi), homogenized in TRIzol reagent for total RNA extraction and viral RNA levels were quantified by RT-qPCR. (**F**) Schematic of the human STAT2 knock-in (hSTAT2-KI) mouse model, in which the endogenous mouse Stat2 locus is replaced with human STAT2. Three-week-old hSTAT2-KI mice (n = 10 per group) were inoculated via footpad injection with WT ZIKV or ZIKV-NS5^L162G^ (1 × 10^4^ FFU per mouse) or mock-injected with PBS. (**G**) Body weight of hSTAT2-KI mice was monitored daily for 7 days after infection with WT ZIKV, ZIKV-NS5^L162G^, or mock infection. (**H-I**) Blood and indicated tissues (brain, heart and spleen) were collected at 4 and 7 dpi from infected or mock-infected hSTAT2-KI mice, homogenized in TRIzol reagent and subjected to total RNA extraction. Viral RNA levels were quantified by RT-qPCR. Bars represent mean ± SEM. Statistical significance was determined using one-way ANOVA with Tukey’s post-hoc test (*p < 0.05; **p < 0.01; ***p < 0.001; ns, not significant).

While A129 mice provide a useful immunodeficient system for assessing viral infection and lethality, the lack of type I IFN signaling precludes evaluation of viral immune evasion mechanisms and may not fully recapitulate disease progression in immunocompetent hosts (35, 51). In contrast, WT mice are largely resistant to ZIKV infection, limiting their applicability for *in vivo* pathogenesis studies (52). To overcome these limitations and enable productive infection under immunocompetent conditions, human STAT2 knock-in (hSTAT2-KI) mice were developed (35, 53, 54) (**Fig. 4F**). These mice are susceptible to ZIKV infection and provide a physiologically relevant model to dissect the role of NS5 in immune evasion and pathogenesis *in vivo*. To evaluate the impact of the L162G mutation, hSTAT2-KI mice were inoculated subcutaneously with WT ZIKV or ZIKV-NS5^L162G^ virus. Body weight was monitored daily and viral RNA loads were quantified in organs and blood at indicated time points (**Fig. 4F**). Whereas WT ZIKV infection caused significant weight loss starting on day 5 post-infection, ZIKV-NS5^L162G^ infected mice maintained body weight similar to mock controls (**Fig. 4G**). Consistently, viral RNA levels in the brain and heart were markedly lower in NS5^L162G^-infected mice than in WT-infected mice (**Fig. 4H**). In addition, viral RNA in the blood was significantly reduced in ZIKV NS5^L162G^-infected hSTAT2-KI mice at 1dpi compared with WT ZIKV infection (**Fig. 4I**), indicating impaired systemic viral dissemination when STAT2 degradation is disrupted. Importantly, viral RNA sequencing from brain tissues of ZIKV-NS5^L162G^ infected hSTAT2-KI mice at 7 dpi confirmed retention of the L162G substitution without reversion (**Fig. S5**), indicating *in vivo* genetic stability.

Collectively, these findings demonstrate that the L162G mutation in NS5 selectively reduces ZIKV infection and disease severity in an immunocompetent host while leaving viral replication intact in an IFN-deficient environment.

### ZIKV-NS5^L162G^ recombinant virus elicits robust cellular and humoral immune responses in vivo

Given that the ZIKV-NS5^L162G^ mutant is attenuated in immunocompetent hSTAT2-KI mice (**Fig.4F-I**), we next investigated whether it remains immunogenic *in vivo*. hSTAT2-KI mice were infected with WT ZIKV or ZIKV-NS5^L162G^, and tissues were collected at 7 and 14 dpi to assess virus-specific T cell responses and cytokine production. Serum samples were collected at 22 dpi to measure neutralizing antibody responses **(Fig. 5A)**.

**Figure 5.**
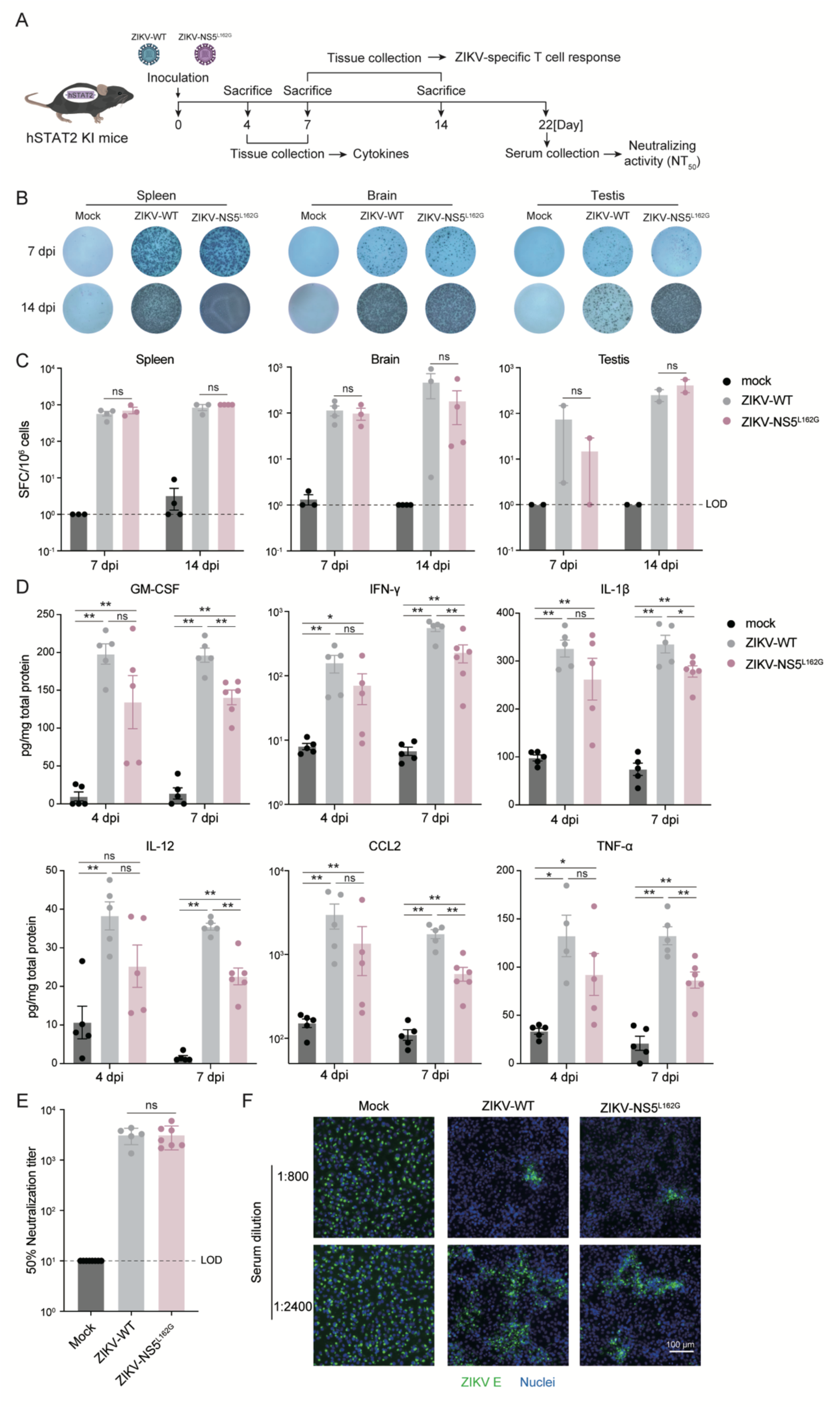
ZIKV-NS5^L162G^ recombinant virus elicits robust cellular and humoral immune responses *in vivo*. (**A**) Schematic of the experimental design. Three-week-old hSTAT2-KI mice were infected via footpad injection with WT ZIKV or ZIKV-NS5^L162G^ (1 × 10^5^ FFU per mouse) or mock infected (n = 25 for mock, n = 22 for WT ZIKV and n = 25 for ZIKV-NS5^L162G^). Spleen, brain and testis were collected at 7 and 14 dpi for analysis of viral E antigen-specific T cell responses (n = 7 for mock, n = 7 for WT ZIKV and n = 7 for ZIKV-NS5^L162G^). Brain tissues were collected at 4 and 7 dpi for cytokine measurements (n = 10 for mock, n = 10 for WT ZIKV and n = 11 for ZIKV-NS5^L162G^). Serum samples were collected at 22 dpi for neutralizing assays (n = 8 for mock, n = 5 for WT ZIKV and n = 7 for ZIKV-NS5^L162G^). (**B-C**) Single-cell suspensions prepared from spleen, brain and testis of mock, WT ZIKV, or ZIKV-NS5^L162G^ infected mice were stimulated *ex vivo* with a synthesized ZIKV E protein peptide library. IFN-γ-secreting T cells were detected by ELISpot. Representative ELISpot images are shown in (B). Quantification of IFN-γ-producing T cells is shown as spot-forming cells (SFC) per 10^6^ cells from the indicated tissues (**C**). (**D**) Brain tissues collected at 4 and 7 dpi from mock, WT ZIKV and ZIKV-NS5^L162G^-infected hSTAT2-KI mice were homogenized and levels of GM-CSF, IFN-γ, IL-1β, IL-12, CCL2 and TNF-α in clarified supernatants were quantified by ELISA and normalized to total protein content. (**E-F**) Serum samples collected at 22 dpi from mock, WT ZIKV and ZIKV-NS5^L162G^-infected hSTAT2-KI mice were analyzed for neutralizing activity by focus reduction neutralization test (FRNT) on Vero cells. The reduction in viral foci was used to determine the 50% neutralization titer (NT_50_). Individual NT_50_ values are shown for each mouse (**E**). Representative immunofluorescence images showing ZIKV neutralization by serially diluted serum samples are shown in (**F**). ZIKV E protein (green) and nuclei (blue) are indicated. Scale bar, 100 μm. Bars represent mean ± SEM from biological replicates. Statistical significance was determined using one-way ANOVA with Tukey’s post-hoc test (*p < 0.05; **p < 0.01; ***p < 0.001; ns, not significant).

To assess ZIKV-specific T cell responses, immune cells were isolated from spleen, brain, and testis at 7 and 14 dpi, stimulated *ex vivo* with a synthesized ZIKV E peptide library (55), and analyzed by IFN-γ ELISpot. ZIKV E-specific IFN-γ responses were detected from both ZIKV WT and NS5^L162G^-infected mice across these tissues compared with mock controls (**Fig. 5B-C**), indicating that the ZIKV-NS5^L162G^ virus efficiently primes antigen-specific T cells despite its reduced pathogenicity. We further measured local inflammatory/antiviral cytokine production in the brain. Brain tissues collected at 4 and 7 dpi were homogenized, and cytokines in clarified supernatants were quantified by ELISA. WT ZIKV infection induced robust increases in multiple cytokines and chemokines, including GM-CSF, IFN-γ, IL-1β, IL-12, CCL2 and TNF-α. Cytokine levels in NS5^L162G^-infected mice were comparable to WT ZIKV infection at 4 dpi but were generally lower at 7 dpi (**Fig. 5D**), suggesting that the ZIKV-NS5^L162G^ mutant triggers effective but less inflammatory immune activation. We further evaluated humoral immunity by measuring serum neutralizing antibody responses at 22 dpi. ZIKV-NS5^L162G^ infected mice developed robust neutralizing antibody responses (NT_50_=3160.86±1453.37), with titers comparable to those elicited by WT ZIKV infection (NT_50_=3147.20 ± 1009.81) (**Fig. 5E-F**), indicating that attenuation of viral replication does not compromise the induction of potent neutralizing humoral immunity.

Together, these data indicate that ZIKV-NS5^L162G^ remains highly immunogenic *in vivo*, inducing both tissue T cell responses and potent neutralizing antibodies while exhibiting reduced pathogenicity.

### Prior exposure to ZIKV-NS5^L162G^ virus confers protection against WT ZIKV challenge

Given that the ZIKV-NS5^L162G^ recombinant virus exhibits reduced pathogenicity while retaining immunogenicity *in vivo*, we next assessed whether prior exposure to this attenuated virus could protect mice from subsequent WT ZIKV challenge. To this end, hSTAT2-KI mice were primed by subcutaneous inoculation with ZIKV-NS5^L162G^ (Dakar-41525 strain, 1 × 10^4^ FFU per mouse) or mock treated. Ten days later, both groups were challenged with WT ZIKV (Dakar-41525 strain, 5 × 10^6^ FFU per mouse), while an additional mock-primed and mock-challenged group served as uninfected controls. Body weight was monitored daily and viral RNA levels in blood and tissues were quantified at the indicated time points (**Fig. 6A**).

**Figure 6.**
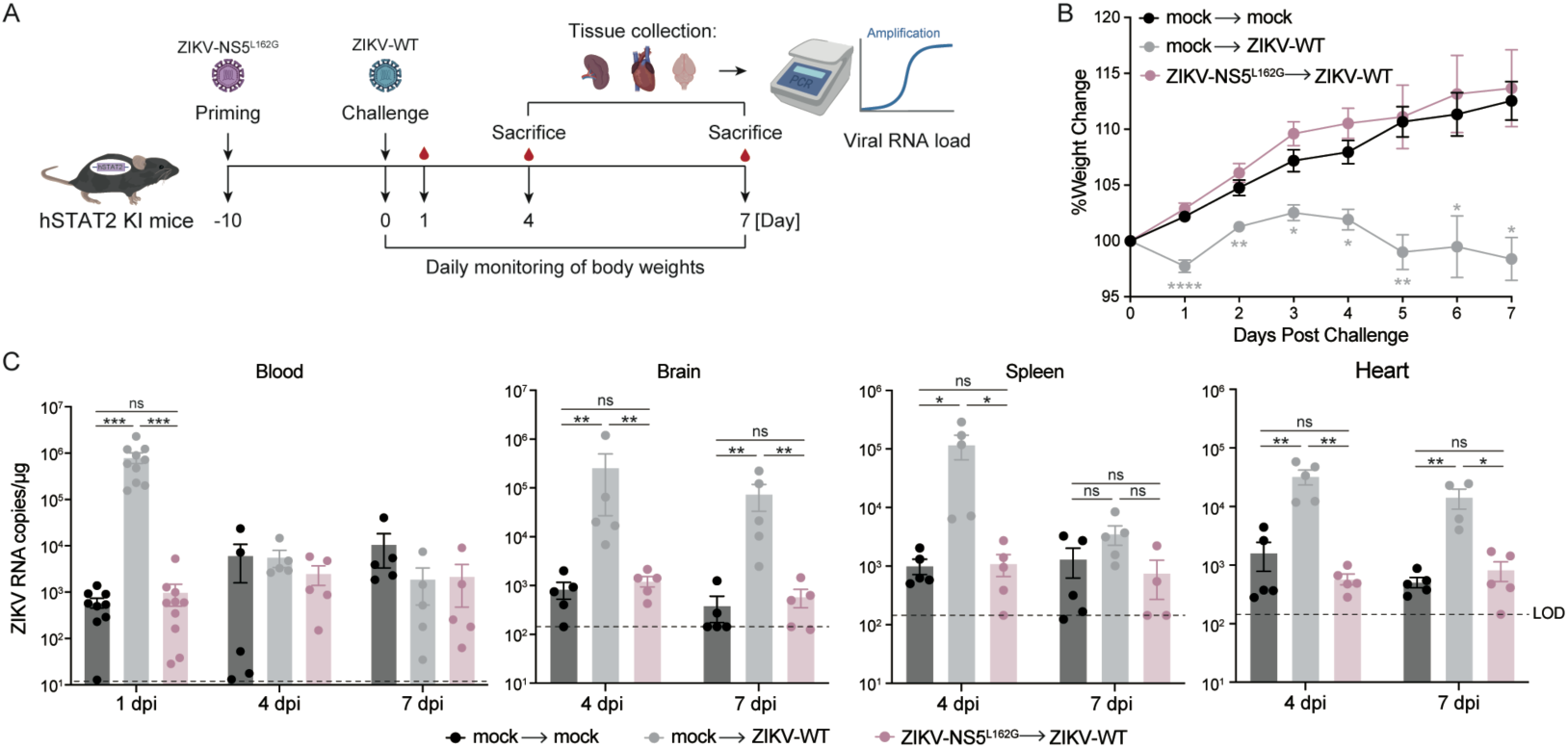
Prior exposure to ZIKV-NS5^L162G^ recombinant virus limits disease and viral replication upon WT ZIKV challenge in hSTAT2-KI mice. **(A)** Schematic of the immunization-challenge experiment. Three-week-old hSTAT2-KI mice were first immunized via footpad injection with ZIKV-NS5^L162G^ (1 × 10^4^ FFU per mouse, n = 10) or mock treated with PBS (n = 20). Ten days later, all ZIKV-NS5^L162G^ exposed mice and ten of the mock-treated mice were challenged with WT ZIKV (Dakar strain, 5 × 10^6^ FFU per mouse), while the remaining ten mock-treated mice received a mock challenge and served as uninfected controls. Body weight was monitored daily after challenge and blood, brain, spleen and heart were collected at the indicated time points for viral RNA quantification. (**B**) Body weight changes of mock-treated, WT ZIKV-challenged mice (gray), ZIKV-NS5^L162G^-exposed and WT ZIKV-challenged mice (pink), and mock-treated, mock-challenged controls (black) were monitored for 7 days post-challenge. (**C**) Blood samples were taken at 1, 4 and 7 dpi, while tissues (spleen, heart, and brain) were collected at 4 and 7 dpi. All samples were homogenized in TRIzol reagent for total RNA extraction and viral RNA levels were quantified by RT-qPCR and expressed as viral RNA copies per μg of total RNA. Bars represent mean ± SEM from biological replicates. Statistical significance was determined using one-way ANOVA with Tukey’s post-hoc test (*p < 0.05; **p < 0.01; ***p < 0.001; ns, not significant).

Following WT ZIKV challenge, mock-primed mice exhibited pronounced weight loss (**Fig. 6B, gray line**). In contrast, mice previously exposed to ZIKV-NS5^L162G^ showed body weight changes that closely overlapped with those of mock-challenged, uninfected controls (**Fig. 6B, pink and black lines**), indicating effective protection. Consistent with these observations, mock-primed mice displayed high levels of viral RNA in the blood, spleen, heart and brain after WT challenge. By contrast, viral RNA in all examined compartments of ZIKV-NS5^L162G^ primed mice remained at or near the detection limit and comparable to that observed in mock-challenged controls (**Fig. 6C**), indicating significant suppression of viral infection following challenge.

Together, these results demonstrate that prior exposure to the STAT2-antagonism-defective ZIKV-NS5^L162G^ virus elicits sterilizing immunity that protects hSTAT2-KI mice from subsequent WT ZIKV infection. This finding provides evidence that targeting NS5-mediated immune evasion can generate a highly attenuated yet protective viral phenotype.

## Discussion

Innate immune antagonism is a hallmark of flavivirus biology, yet the physiological contribution of individual evasion strategies to viral fitness and disease remains incompletely defined. In this study, we decoupled the replication and immune-antagonistic functions of ZIKV NS5 through a multistep functional screening strategy that jointly assessed STAT2 degradation and viral RNA replication. By combining IFN signaling-based reporter assays, a STAT2 reporter cell system to quantitatively monitor STAT2 abundance, and replicon-based replication profiling (**Fig.1; Fig S1-S2**), we identified residue L162 of NS5 as uniquely permissive to a set of substitutions that abolish NS5-mediated STAT2 degradation while preserving viral RNA replication (**Fig. 1B-D**). Using reverse genetics, we generated recombinant viruses bearing these mutations (L162A or L162G) and demonstrated that they fail to recruit the ZSWIM8-CUL3 E3 ligase, cannot promote STAT2 proteasomal turnover (**Fig. 2**), and consequently restore IFN signaling and ISG induction in human cells (**Fig. 3**). Both *in vitro* and *in vivo* analyses further establish that these viruses are restricted in a STAT2-dependent manner, establishing that NS5-mediated STAT2 degradation is a critical determinant of ZIKV replication fitness and pathogenesis in an IFN-competent host (**Fig. 4**). Notably, despite their attenuated infection phenotype, NS5 L162 mutant viruses retain robust immunogenicity *in vivo* and prior exposure markedly limits viral replication and disease upon subsequent WT challenge (**Fig. 5-6**), underscoring the functional importance of NS5-mediated STAT2 antagonism in shaping viral fitness, pathogenesis and host immunity.

Our findings refine the current understanding of NS5 as a multifunctional protein integrating viral RNA synthesis with immune modulation. Veit et al. provided evolutionary evidence that NS5-mediated antagonism of STAT2 is a conserved strategy among vector-borne flaviviruses, with resistance or susceptibility emerging repeatedly across mammalian lineages (56), and In line with this, NS5 from ZIKV and DENV suppresses type I IFN signaling through species-specific targeting of STAT2 (32, 56, 57). Indeed, WT mice are resistant to ZIKV and DENV infection because murine STAT2 is refractory to NS5 antagonism, and productive infection requires either genetic ablation of type I IFN signaling or replacement of murine STAT2 with the human ortholog (34, 35, 58). We found that these two NS5 functions are genetically inseparable at nearly all positions examined. Strikingly, residue L162 emerged as the sole site at which specific substitutions selectively disrupt STAT2 degradation while leaving viral replication intact (**Fig. 1**). Structural analyses of the reported ZIKV NS5-STAT2 complex place residue L162 within the NS5 methyltransferase domain and outside the STAT2 interaction interface (**Fig. 2D**), consistent with our biochemical data showing that L162 substitutions do not affect NS5-STAT2 binding (**Fig. 2C**). In contrast, these substitutions selectively disrupt NS5 interaction with ZSWIM8, suggesting that L162 contributes to a distinct interaction surface required for recruitment of the ZSWIM8-CUL3 E3 ligase complex. Although structural information for the NS5-ZSWIM8-CUL3 complex is currently unavailable, our data support a model in which L162 specifically mediates ligase engagement rather than STAT2 recognition, thereby functionally uncoupling immune antagonism from replication activity. This distinction underscores the power of unbiased functional screening approaches to reveal separable regulatory nodes within multifunctional viral proteins that may not be apparent from structural or candidate-based analyses alone.

Beyond its mechanistic insights, our study has broader translational implications. First, it provides *in vivo* validation that preservation of STAT2 function has a meaningful antiviral effect, supporting the NS5-ZSWIM8-CUL3-STAT2 axis as a tractable target for host-directed antiviral strategies. The small molecule (e.g., SMU-1k) that blocks NS5-mediated STAT2 degradation have recently been reported to restore IFN signaling and reduce viral cytopathic effects in cell culture (59). Our findings extend these observations by demonstrating the physiological relevance of this pathway *in vivo*, underscoring the therapeutic potential of targeting viral immune antagonism. Second, although our study was not designed as a vaccine development effort, it highlights a general principle with potential relevance to attenuation strategies. The ZIKV-NS5^L162G^ mutant virus retains replication competence in IFN-deficient settings yet is markedly restricted in IFN-competent hosts (**Fig.4**), while still eliciting robust cellular and humoral immune responses and conferring protection against subsequent WT challenge *in vivo* (**Fig. 5-6**). These findings provide proof-of-principle that selective disruption of viral immune antagonism can yield attenuated yet immunogenic viral phenotypes without directly compromising intrinsic replication machinery.

Several important considerations remain. Because attenuation in this context depends on an intact type I IFN response, safety in immunocompromised hosts is a concern, as illustrated by the retained pathogenicity of the mutant virus in IFN receptor-deficient mice (**Fig. 4A-E**). In humans, a subset of individuals harbor inborn errors of type I IFN immunity or neutralizing autoantibodies against type I IFNs, which have been shown to confer heightened susceptibility to severe viral diseases, including COVID-19 and influenza (60–62). In such settings, attenuation based solely on impaired IFN antagonism may be insufficient, and viruses lacking NS5-mediated STAT2 degradation could still retain substantial pathogenic potential. A rational strategy to mitigate this limitation is to combine immune-evasion-targeted attenuation with additional, mechanistically independent attenuating changes. For example, coupling disruption of NS5-mediated STAT2 degradation with mutations that impair other flaviviral immune antagonists could reduce reliance on a single host pathway for attenuation and substantially lower the risk of pathogenicity in IFN-compromised settings. Such combinatorial attenuation strategies would also decrease the likelihood of genetic reversion while preserving sufficient replication competence to elicit durable immune responses. More broadly, our findings establish immune-evasion nodes as tunable determinants of viral fitness and provide a conceptual framework for extending this approach to other flaviviruses. Applying similar strategies to viruses such as DENV, which exploit distinct E3 ligase (UBR4) complexes to antagonize STAT2 (30), may further clarify conserved and divergent roles of NS5-mediated immune evasion across the genus and inform the rational balancing of attenuation, immunogenicity, and safety.

In conclusion, by decoupling the replication and immune-antagonistic functions of ZIKV NS5, we demonstrate that NS5-mediated STAT2 degradation is a key driver of immune evasion, viral fitness and pathogenesis *in vivo*. This work not only advances fundamental understanding of flavivirus-host interactions but also establishes a conceptual and experimental framework for targeting viral immune antagonism in the development of host-directed antivirals and rational attenuation strategies.

## Acknowledgement

We thank Dr. Adolfo Garcia-Sastre (Icahn School of Medicine at Mount Sinai, US) for providing the human STAT2 cDNA clone, and Dr. Chao Shan (Wuhan Institute of Virology, CAS, China) for sharing the pACYC-Dakar-41525 ZIKV infectious clone. We are grateful to Dr. Gang Long (Fudan University) for generously providing the pFK-ZIKA(KU321639)-Gluc replicon plasmid and NS5 antibody, and to Dr. Gong Cheng (Tsinghua University) for providing A129 mice. We thank Drs. Xiaohui Ju, Mingli Gong and Qingqing Li (Ding laboratory alumni) for their contributions to the NS5 mutant panel. We also thank all members of the Ding laboratory for insightful discussions and critical comments on the manuscript. Technical support from the Laboratory Animal Resources Center at Tsinghua University is sincerely appreciated.

This work was supported by the National Key Research and Development Plan of China (2023YFC2305900), National Natural Science Foundation of China (82241077, 82341084 and 82272302), Tsinghua University Dushi Program (20251080029) and SXMU-Tsinghua Collaborative Innovation Center for Frontier Medicine. The funders had no role in study design, data collection and analysis, decision to publish, or preparation of the manuscript.

## Materials and Methods

### Cell culture

HEK293T (ATCC, CRL-3216), A549 (ATCC #CCL-185), and Vero E6 (Cell Bank of the Chinese Academy of Sciences, Shanghai, China) cells were maintained in Dulbecco’s modified Eagle medium (DMEM) (Gibco, NY, USA) supplemented with 10% (vol/vol) fetal bovine serum (FBS), 10mM HEPES, 1mM sodium pyruvate, 1×non-essential amino acids and 50 IU/ml penicillin/streptomycin. All cells were cultured at 37°C in a humidified atmosphere containing 5% CO₂. Cell lines were routinely tested and confirmed to be free of mycoplasma contamination.

### Plasmids

The cDNAs encoding human STAT2, WT ZIKV NS5 and NS5 mutants were cloned into the pLVX-IRES-zsGreen1 expression vectors (Catalog No. 632187, Clontech Laboratories, Inc) using standard molecular cloning techniques. All of the constructs were verified by Sanger sequencing.

### Reporter assays

HEK293T cells (2×10^5^ per well) were seeded in 24-well plates and co-transfected with 500 ng of plasmids expressing wild-type or mutant NS5 proteins, along with 100 ng of the firefly luciferase reporter plasmid pGL4-ISRE (driven by the interferon-stimulated response element) and 10 ng of the Renilla luciferase control plasmid pRL-TK, using Vigofect (Vigorous Biotechnology). After 24 h, cells were stimulated with human IFN-β (10 ng/mL) for an additional 24 hours. Cells were lysed, and luciferase activity was measured using the Dual-Luciferase Reporter Assay System (Promega, Madison, WI, USA) according to the manufacturer’s instructions. The firefly luciferase activities were normalized to Renilla luciferase activities.

### Western blotting assay

Western blotting was performed as previously described (63). Briefly, protein samples were separated by SDS-PAGE and transferred onto PVDF membranes. Membranes were blocked with 5% nonfat milk in 1×PBS containing 0.1% Tween-20 (PBST) for 1 h at room temperature and then incubated for 2 h with primary antibodies diluted in the same blocking buffer. The following primary antibodies were used: anti-Flag (Lablead, #F005), anti-HA (ABclonal, #AE008, anti-STAT2 (Sangon, #D261445), anti-ZIKV NS5 (a gift from Dr. Gang Long at Fudan University), β-actin (Abcepta, #AM1021b), and anti-tubulin (CWBIO, #CW0098). After three washes in PBST, membranes were incubated for 1 h with HRP-conjugated secondary antibodies (Abclonal, #AS003 and #AS014), followed by additional washes. Protein bands were visualized using enhanced chemiluminescence (ECL) and imaged with an ImageQuant LAS 4000 luminescent analyzer (GE Healthcare).

### Immunoprecipitation assay

For each sample, about 2×10^6^ cells were washed with ice-cold PBS and lysed in 400 μl Protein Lysis Buffer (same as WB assay) on ice for 30 min. Lysates were centrifuged at 12000 rpm at 4°C for 15 min, and supernatant was transferred to new tubes. 50 μL was saved as input sample. For immunoprecipitation of HA-tagged proteins, lysates were incubated with 5 μl Anti-HA-Nanoab-Magnetic beads (Lablead) for 6 h at 4°C while rotating. Beads were washed 5 times with lysis buffer beads were mixed with 120 μL 1×SDS-loading buffer.

### *In vitro* transcription

PCR-assembled DNA fragments (for ZIKV replicon) or linearized plasmid DNA (for recombinant virus genomes) were purified by phenol/chloroform extraction followed by isopropanol precipitation. Approximately 1 μg of purified DNA was used as a template for *in vitro* transcription with the EasyCap T7 Co-transcription Kit with CAG Trimer (Vazyme, China), following the manufacturer’s protocol. The resulting capped RNA transcripts were purified by LiCl precipitation, resuspended in nuclease-free water, and quantified spectrophotometrically. RNA integrity was verified by agarose gel electrophoresis.

### ZIKV replicon assay

The pFK-ZIKA (KU321639)-Gluc replicon plasmid lacking the T7 promoter was used as the template to amplify fragment A and fragment B (Fig. 2E). During amplification of fragment A, the T7 promoter sequence and the desired NS5 point mutations were introduced simultaneously. The full-length replicon DNA fragment was then assembled by overlap-extension PCR using PrimeSTAR® GXL DNA Polymerase (Takara, Japan). The assembled DNA products (1 μg each) were used as templates for *in vitro* transcription as described above. For RNA transfection, 500 ng of *in vitro*-transcribed RNA was transfected into approximately 2×10^5^ cells using the TransIT-mRNA Transfection Kit (Mirusbio) following the manufacturer’s instructions. Following transfection, cells were washed three times with DPBS at 8 h, 24 h and 48 h post-transfection to remove residual input RNA. Culture supernatants were collected at 72 h post-transfection and Gaussia luciferase activity was measured using the Renilla Luciferase Assay System (Promega, #E2820). Luminescence was recorded on a GloMax luminometer (Promega).

### Recombinant ZIKV production, titration and infection assays

WT ZIKV and NS5 mutant viruses (NS5^L162A^ and NS5^L162G^) were generated using the pACYC-Dakar-41525 infectious clone backbone. Infectious clone plasmids were linearized with ClaI (NEB), followed by phenol-chloroform extraction and ethanol precipitation. Linearized DNA (1 μg) was used as template for *in vitro* transcription as described above. For virus rescue, 10 μg of *in vitro*-transcribed RNA was electroporated into 8×10^6^ Vero cells using the Gene Pulser Xcell™ Electroporation Systems (Bio-rad) at 270 V and 950 μF. Electroporated cells were transferred to culture flasks and incubated at 37°C. Supernatants were harvested 48 h after electroporation, clarified, aliquoted and stored at −80 °C as recombinant virus stocks. Viral titers were determined by focus-forming assay (FFA) on Vero cells. Briefly, 10-fold serial dilutions of virus-containing supernatants were added to confluent Vero monolayers in 24-well plates for 1 h at 37 °C. After adsorption, cells were overlaid with DMEM containing 1% carboxymethyl cellulose and incubated for 48 h. Monolayers were fixed with 4% paraformaldehyde, permeabilized and stained with the flavivirus-cross-reactive monoclonal antibody 4G2 followed by an HRP-conjugated secondary antibody. Foci were visualized using TrueBlue Peroxidase Substrate (KPL) and counted to calculate viral titers (FFU/ml). For infection assays, Vero or A549 cells were infected with WT or NS5 mutant viruses at the indicated multiplicity of infection (MOI). After 1 h of viral adsorption at 37 °C, inocula were removed, cells were washed with PBS and fresh medium was added. Infected cells were monitored for cytopathic effect (CPE) and harvested at the indicated time points for immunofluorescence staining, RNA extraction, or other downstream analyses.

### RNA-seq and data analysis

A549 cells were infected with WT ZIKV, ZIKV-NS5^L162G^, ZIKV-NS5^L162A^ (moi=0.05), or mock treated. After 48h, total RNA was extracted and submitted to the Annoroad Gene Technology (Beijing) for RNA library preparation and high-throughput sequencing using the DNBSEQ T7 platform. Clean reads were aligned to the human reference genome (Homo_sapiens GRCh38.p13) using HISAT2 (v2.1.0). Gene expression levels were quantified with RSEM (v1.3.1) and differential expression analysis was performed using DESeq2. Genes with p < 0.05 and |log_2_ fold change| ≥ 1 were considered differentially expressed (DEGs).

### Mice

C57BL/6 hSTAT2 KI mice were purchased from the Jackson Laboratory (C57BL/6-*Stat2^tm1.1(STAT2)Diam^*/AgsaJ, strain #:031630). A129 mice (IFNAR^-/-^) on a C57BL/6 genetic background were kindly provided by Dr. Gong Cheng (Tsinghua University). All mice were bred in the Laboratory Animal Resources Center of Tsinghua University. All animal experiments were performed in accordance to a protocol (number: 24-DQ3) reviewed and approved by the Institution Animal Care and Use Committee (IACUC) of Tsinghua University.

### Mice infection, body weight monitoring and RNA extraction from serum and tissues

A129 or hSTAT2-KI mice were infected via footpad injection with ZIKV diluted in 20 μL of NTE buffer. Mock-infected controls received an equivalent volume of NTE buffer alone. Body weight was monitored daily and mortality was recorded throughout the course of infection. At the indicated time points, mice were anesthetized with isoflurane and approximately 20 μL of blood was collected via retro-orbital bleeding into 1.5 mL microcentrifuge tubes. Viral RNA was extracted from whole blood using the RaPure Total RNA Kit (Magen, China) according to the manufacturer’s instructions. At indicated time points, mice were euthanized by carbon dioxide inhalation, and the spleen, brain and heart were collected. Approximately 30-40 mg of each tissue was homogenized in TRIzol reagent (Invitrogen, USA) using a mechanical homogenizer and total RNA was isolated following the manufacturer’s protocol.

### ELISpot assay

ZIKV-specific T cell responses were analyzed using an IFN-γ ELISpot assay, as described previously (64). Briefly, 96-well ELISpot plates (Millipore) were coated overnight at 4°C with anti-mouse IFN-γ antibody (BD Biosciences) diluted 1:200 in PBS (100 μL per well). Plates were washed and blocked with RPMI-1640 medium containing 10% fetal bovine serum (FBS) for 2 h at room temperature. Cells isolated from the organs of infected or control mice were seeded for analysis. Splenocytes were seeded at 5 ×10^5^ cells per well, immune cells from the brains and testes were seeded at 1×10^5^ cells per well due to lower cell yields from these tissues, due to lower cell yields. Cells were stimulated with a synthesized ZIKV E protein peptide library (10 μg/mL per peptide), while unstimulated wells (medium only) served as negative controls, and PMA-treated wells were used as positive controls. Cells were incubated at 37°C with 5% CO_2_ for 18h. After incubation, plates were washed with deionized water. Biotinylated anti-mouse IFN-γ detection antibody (BD Biosciences, 1:250 dilution) was added at 100 μL per well and incubated for 2 h at room temperature. Plates were washed three times with PBST, followed by incubation with Streptavidin-HRP (1:500 dilution, 100 μL per well) for 1 h. After washing, spots were developed using TMB substrate for 5-10 min and rinsed thoroughly with distilled water. Plates were air-dried and analyzed using an automated ELISpot reader (ImmunoSpot CTL). The number of IFN-γ-producing cells (spot-forming cells, SFCs) was normalized to 10^6^ cells as the denominator. Data were expressed as mean ± SEM from biological replicates.

### Statistical analysis

Statistical analysis was performed using GraphPad Prism version 8. The data underwent analysis via Student’s unpaired *t* test or ANOVA as applicable. A significance level of P < 0.05 was deemed statistically significant. Results are presented as means ± SEM.

## Supplemental Figure and Figure legends

**Figure S1.**
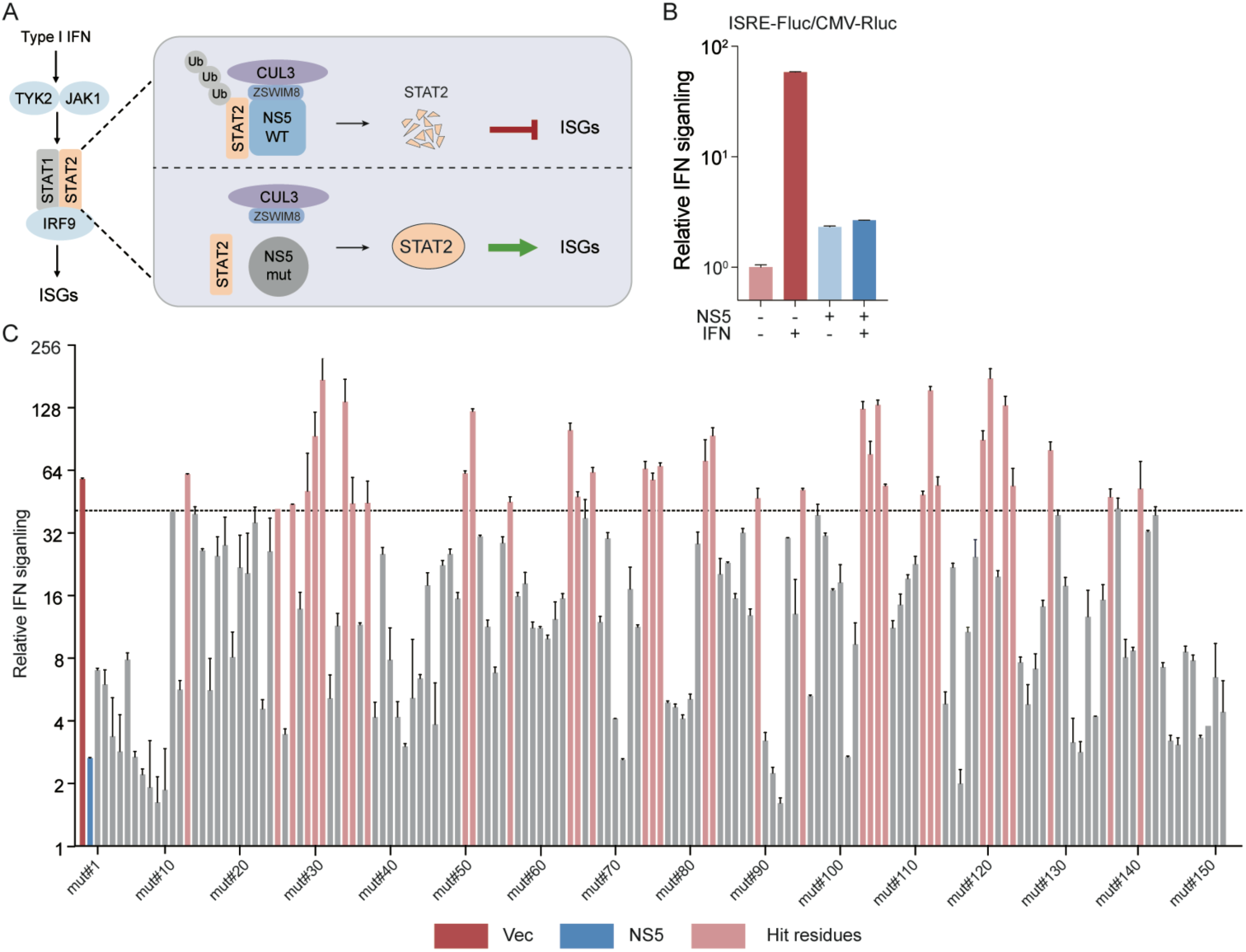
Primary screening for NS5 regions required for IFN antagonism. (**A**) ZIKV NS5 recruits the host ZSWIM8-CUL3 E3 ubiquitin ligase complex to promote STAT2 ubiquitination and proteasomal degradation, thereby suppressing IFN-induced ISG expression. NS5 mutations that disrupt this process are expected to restore IFN signaling and ISG induction. (**B**) Individual NS5 alanine-scanning mutants (151 total; 500 ng each) were transfected in HEK293T cells (2×10^5^ cells per well) together with an ISRE-driven firefly luciferase reporter (100 ng) and a CMV promoter-driven Renilla luciferase construct (10 ng) as an internal control. At 48 h post-transfection, cells were stimulated with IFN for 24 h and luciferase activities were subsequently measured. (**C**) Firefly luciferase activity was normalized to Renilla luciferase and expressed relative to WT NS5 for all 151 NS5 alanine-scanning mutants. Mutants that failed to suppress IFN-induced ISRE activation are highlighted in red.

**Figure S2.**
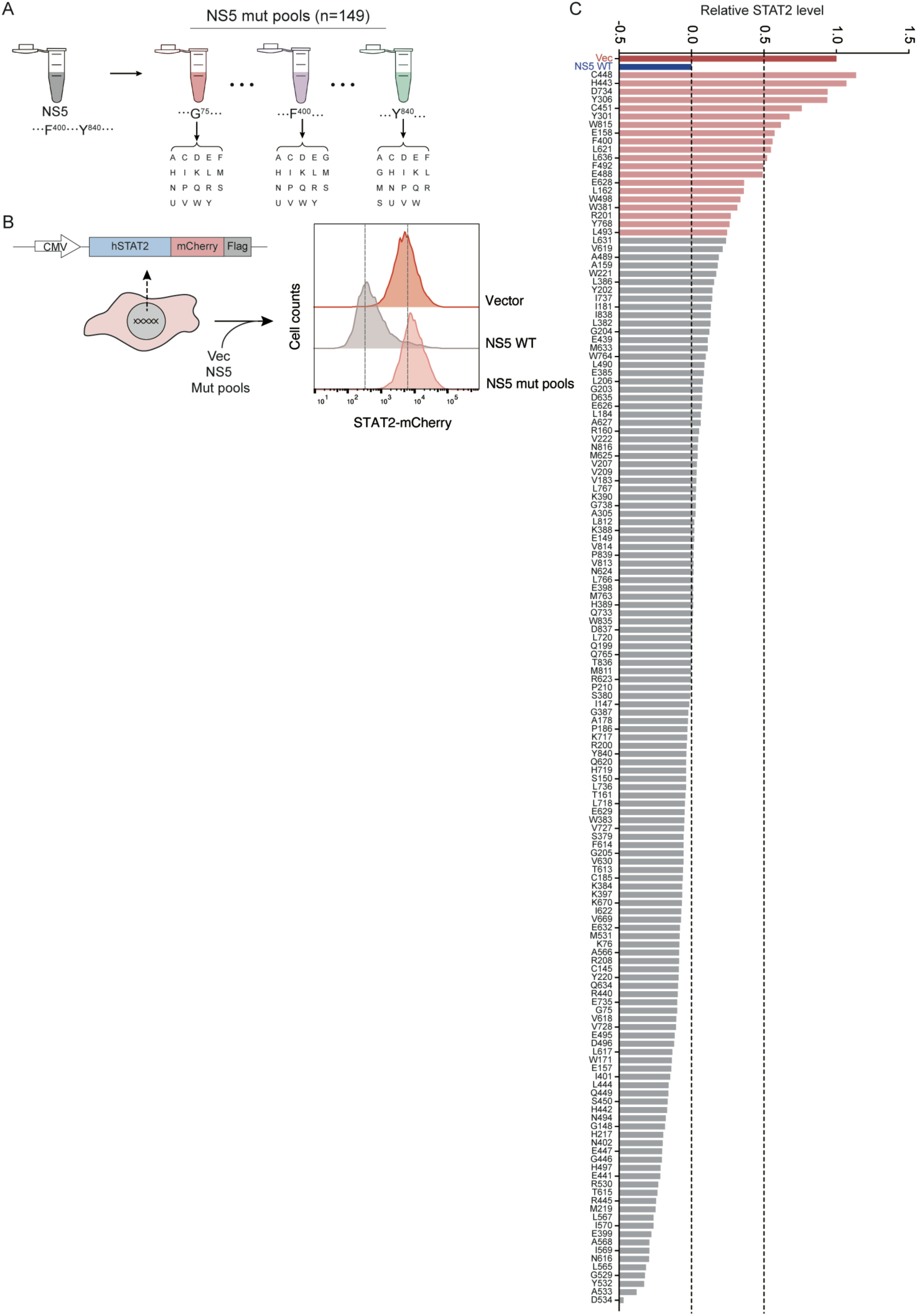
Deep mutational scanning to refine NS5 residues required for STAT2 degradation. (**A-B**) Based on the secondary alanine-scanning screen (**Table S1**), 149 individual NS5 residues were selected for deep mutational analysis. For each residue, a plasmid pool encoding all 19 possible single-amino-acid substitutions was generated (**A**). Each pooled library was individually transfected into STAT2-mCherry reporter cells (2×10^5^ cells per well). At 48 h post-transfection, STAT2 abundance was quantified by flow cytometry based on mCherry fluorescence intensity (**B**). (**C**) Quantification of STAT2-mCherry mean fluorescence intensity (MFI) for each residue-specific mutation pool, normalized to the vector control. Residue pools exhibiting increased STAT2 abundance, indicative of impaired NS5-mediated STAT2 degradation, are highlighted.

**Figure S3.**
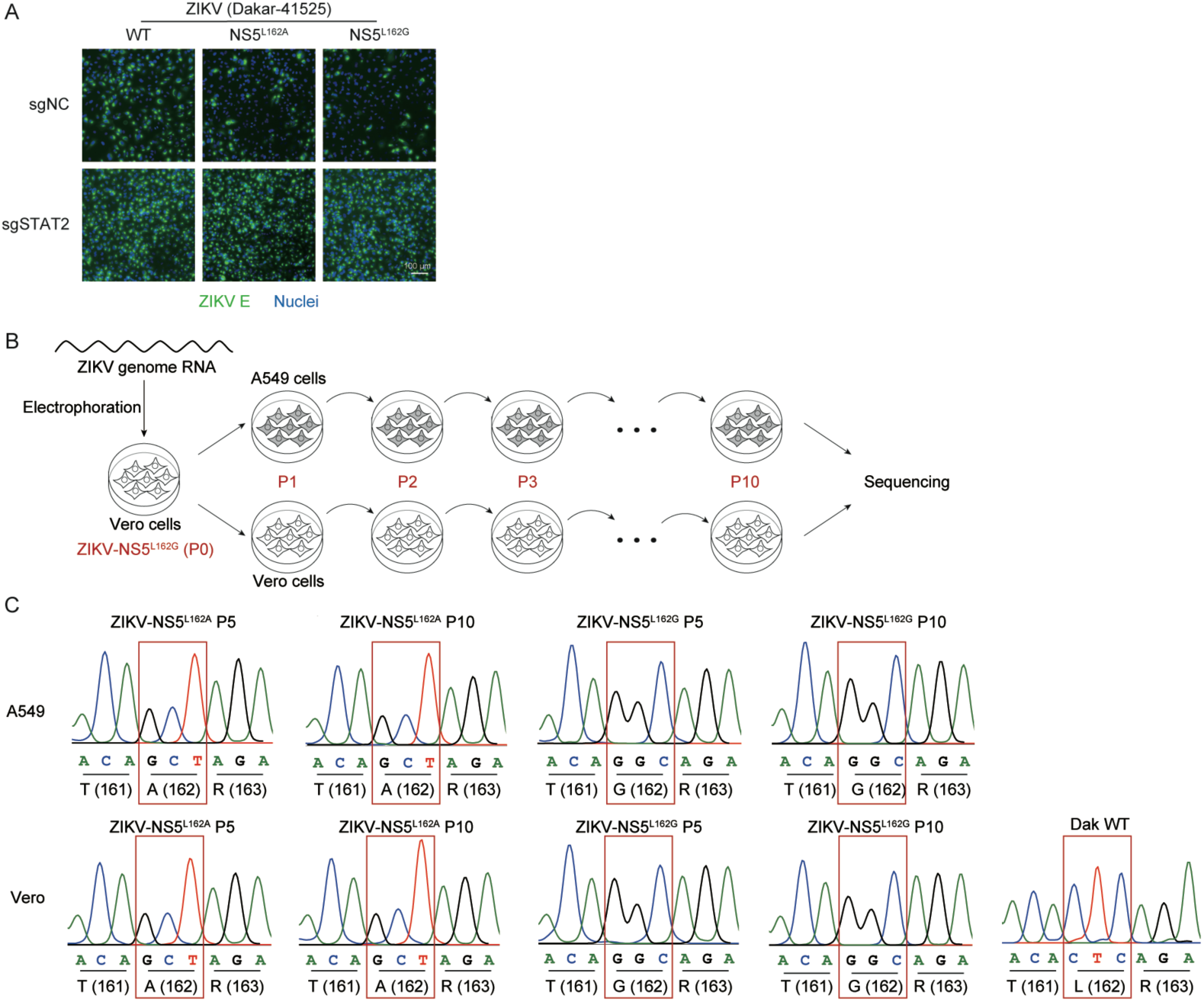
Infection and genetic stability of NS5 mutant viruses in cell culture. (**A**) WT A549 cells (sgNC) or STAT2 knockout A549 cells (sgSTAT2) were infected with WT ZIKV, ZIKV-NS5^L162A^, or ZIKV-NS5^L162G^ at an MOI of 0.01. At 48 h post-infection, cells were fixed with paraformaldehyde, permeabilized and immunostained for ZIKV E protein using the flavivirus-reactive monoclonal antibody 4G2 (green). Nuclei were counterstained with DAPI (blue). Representative images are shown. Scale bar, 100 μm. (**B**) Schematic overview of serial passage experiments. WT ZIKV and NS5 mutant viruses were serially passaged in WT A549 cells or Vero cells for up to 10 passages. At each passage, infected cells were harvested, viral RNA was extracted and the NS5 coding region was amplified by RT-PCR for sequence analysis. (**C**) Representative Sanger sequencing chromatograms of the NS5 region encompassing residue 162 from passages 5 (P5) and 10 (P10) in A549 or Vero cells,

**Figure S4.**
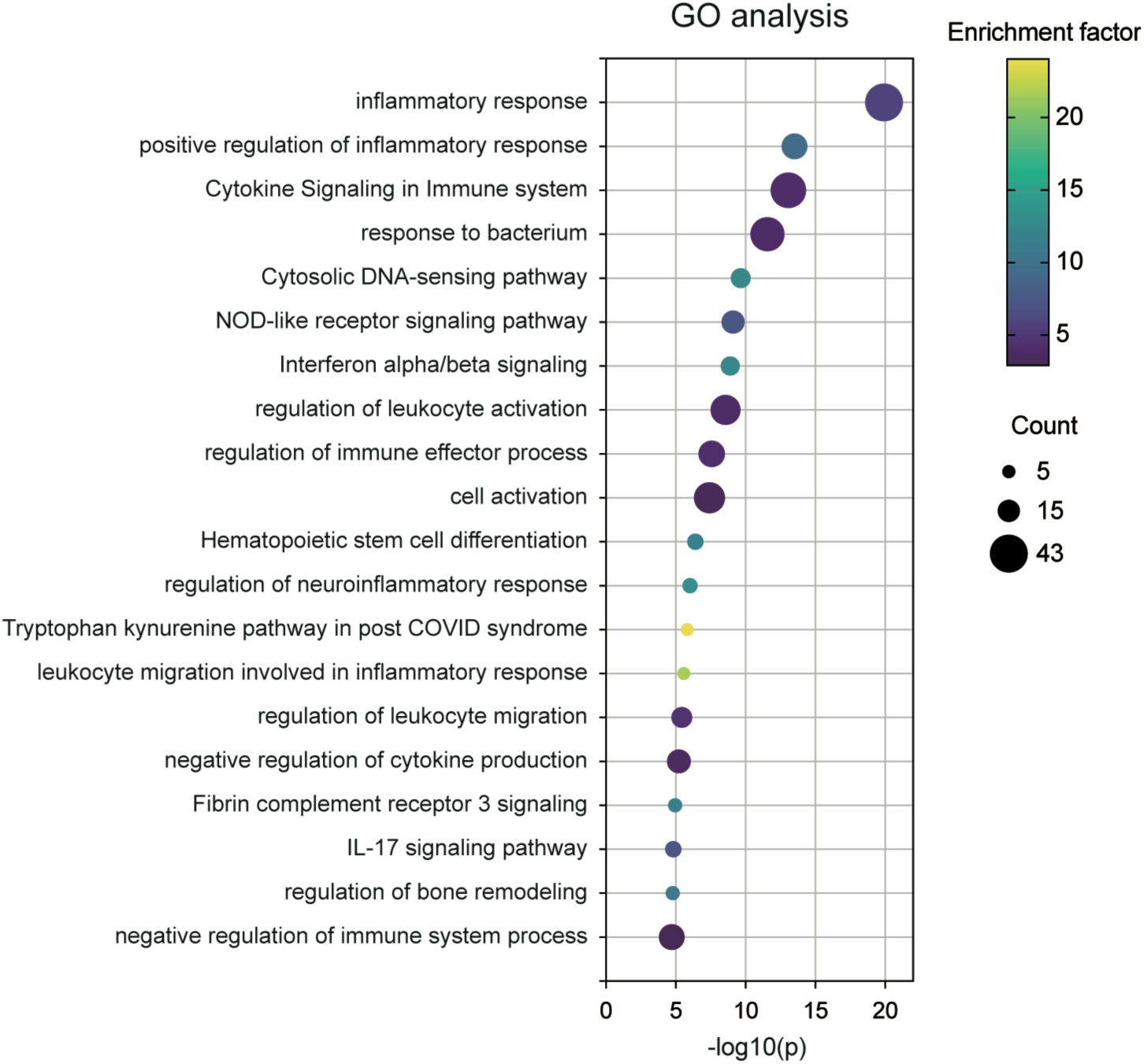
Gene Ontology analysis of host transcriptional responses to NS5 mutant virus infection. Gene Ontology (GO) enrichment analysis of genes upregulated in WT A549 cells infected with ZIKV-NS5^L162A^ or ZIKV-NS5^L162G^ relative to WT ZIKV infection, based on the RNA-seq data shown in Fig. 3D. GO analysis was performed using the Metascape web-based analysis platform. Terms with a p value < 0.01, a minimum gene count of 3 and an enrichment factor > 1.5 were retained and grouped into functional clusters based on membership similarity.

**Figure S5.**
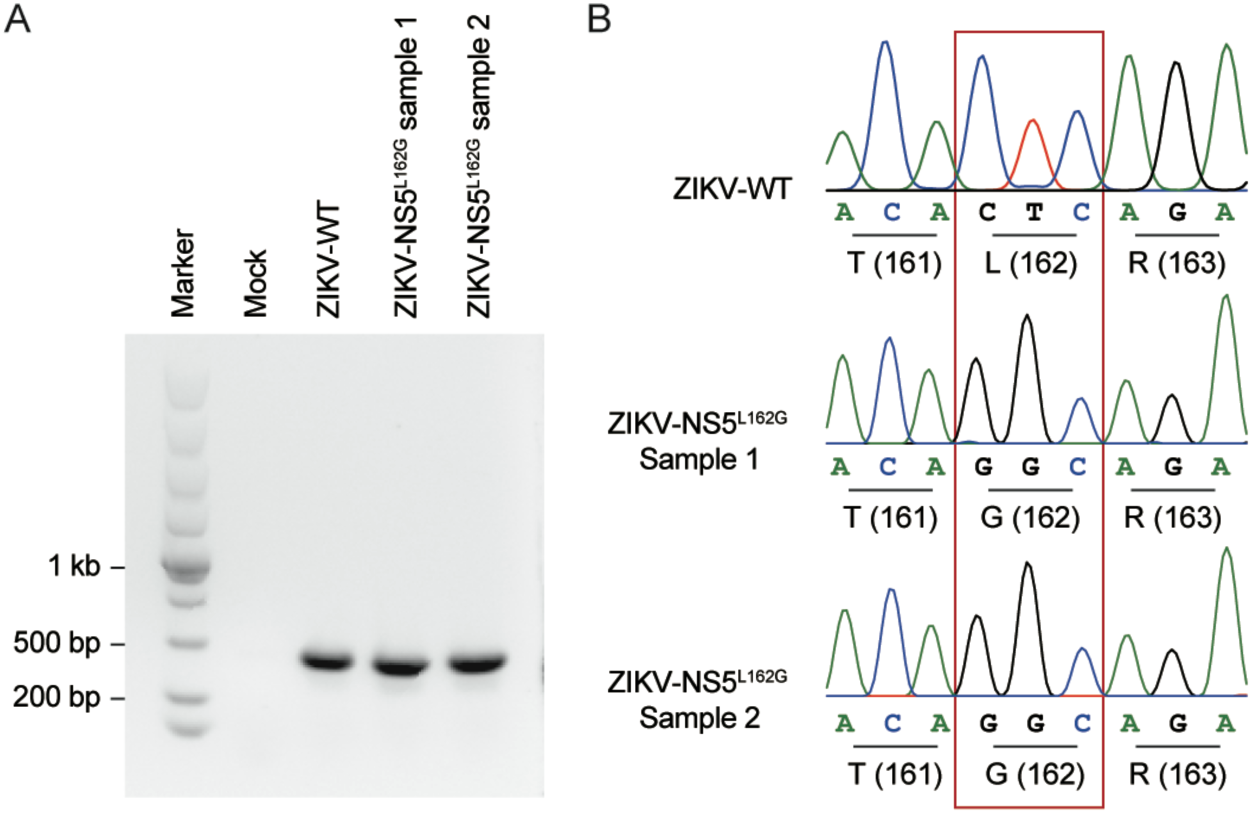
Genetic stability of the ZIKV-NS5L162G mutant virus *in vivo*. (**A**) Viral RNA was isolated from brain tissues of two independent hSTAT2-KI mice infected with ZIKV-NS5L162G at 7 days post-infection. The NS5 coding region encompassing residue 162 was amplified by RT-PCR and subjected to Sanger sequencing. (**B**) Representative sequencing chromatograms spanning the L162 region of NS5 from the two mice. The mutated codon corresponding to L162G is indicated.

## Supplemental Table

**Table S1. Secondary screening for NS5 regions required for STAT2 degradation.**

