## Supplementary material for "A tunable immune-evasion function of Zika virus NS5 governs viral fitness and pathogenesis": Table S1

| Number | | Relative STAT2 level | Site | Original sequence | Mutated sequence |
|---|---|---|---|---|---|
| | vec | 0.91 | | | |
| | NS5 | 0.54 | | | |
| | #13 | 0.96 | | | |
| **#13 d** | mut 1 | 0.85 | 73-78 | PYGKVI | AAAAVI |
| **#13 e** | mut 2 | 0.90 | 73-78 | PYGKVI | AAGKAA |
| **#13 f** | mut 3 | 0.97 | 73-78 | PYGKVI | PYAAAA |
| **#29 d** | mut 4 | 0.96 | 169-174 | GDWLEK | AAAAEK |
| **#29 e** | mut 5 | 0.77 | 169-174 | GDWLEK | AAWLAA |
| **#29 f** | mut 6 | 0.93 | 169-174 | GDWLEK | GDAAAA |
| **#29 GDMAAA** | mut 7 | 0.73 | 169-174 | GDWLEK | GDWAAA |
| **#29 EA** | mut 8 | 0.95 | 169-174 | GDWLEK | GDWLEA |
| **#29 AK** | mut 9 | 0.95 | 169-174 | GDWLEK | GDWLAK |
| **#51 c** | mut 10 | 0.96 | 301-306 | YRTWAY | YRTWAA |
| **#51 d** | mut 11 | 0.98 | 301-306 | YRTWAY | AAAAAY |
| **#51e** | mut 12 | 0.98 | 301-306 | YRTWAY | AATWAA |
| **#51 YAAAAY** | mut 13 | 0.68 | 301-306 | YRTWAY | YAAAAY |
| **#51 ARAAAY** | mut 14 | 0.97 | 301-306 | YRTWAY | ARAAAY |
| **#51 ARAAAA** | mut 15 | 0.98 | 301-306 | YRTWAY | ARAAAA |
| **#51 AW** | mut 16 | 0.96 | 301-306 | YRTWAY | YRAWAY |
| **#51 TA** | mut 17 | 0.96 | 301-306 | YRTWAY | YRTAAY |
| **#50/#51** | mut 18 | 0.96 | 295-306 | FDENHPYRTWAY | AAAAAAAAAAAA |
| **#76 d** | mut 19 | 0.65 | 451-456 | CVYNMM | AAAAMM |
| **#76 e** | mut 20 | 0.96 | 451-456 | CVYNMM | AAYNAA |
| **#76 f** | mut 21 | 0.90 | 451-456 | CVYNMM | CVAAAA |
| **#76 AM** | mut 22 | 0.97 | 451-456 | CVYNMM | CVYNAM |
| **#76 MA** | mut 23 | 0.90 | 451-456 | CVYNMM | CVYNMA |
| **#76 AV** | mut 24 | 0.83 | 451-456 | CVYNMM | AVYNMM |
| **#76 CA** | mut 25 | 0.61 | 451-456 | CVYNMM | CAYNMM |
| **#82 d** | mut 26 | 0.96 | 487-492 | FEALGF | AAAAGF |
| **#82 e** | mut 27 | 0.96 | 487-492 | FEALGF | AAALAA |
| **#82 f** | mut 28 | 0.95 | 487-492 | FEALGF | FEAAAA |
| **#82 AEALAA** | mut 29 | 0.96 | 487-492 | FEALGF | AEALAA |
| **#82FAALAA** | mut 30 | 0.96 | 487-492 | FEALGF | FAALAA |
| **#82 AF** | mut 31 | 0.76 | 487-492 | FEALGF | FEALAF |
| **#82 GA** | mut 32 | 0.95 | 487-492 | FEALGF | FEALGA |
| **#30 d** | mut 33 | 0.58 | 175-180 | RPGAFC | AAAAFC |
| **#30 e** | mut 34 | 0.94 | 175-180 | RPGAFC | AAGAAA |
| **#30 f** | mut 35 | 0.77 | 175-180 | RPGAFC | RPAAAA |
| **#34 d** | mut 36 | 0.82 | 199-204 | QRRYGG | AAAAGG |
| **#34 e** | mut 37 | 0.84 | 199-204 | QRRYGG | AARYAA |
| **#34 f** | mut 38 | 0.96 | 199-204 | QRRYGG | QRAAAA |
| **#50 d** | mut 39 | 0.77 | 295-300 | FDENHP | AAAAHP |
| **#50 e** | mut 40 | 0.92 | 295-300 | FDENHP | AAENAA |
| **#50 f** | mut 41 | 0.58 | 295-300 | FDENHP | FDAAAA |
| **#83 d** | mut 42 | 0.94 | 493-498 | LNEDHW | AAAAHW |
| **#83 e** | mut 43 | 0.95 | 493-498 | LNEDHW | AAEDAA |
| **#83 f** | mut 44 | 0.95 | 493-498 | LNEDHW | LNAAAA |
| **#103 d** | mut 45 | 0.93 | 613-618 | TFTNLV | AAAALV |
| **#103 e** | mut 46 | 0.90 | 613-618 | TFTNLV | AATNAA |
| **#103 f** | mut 47 | 0.86 | 613-618 | TFTNLV | TFAAAA |
| **#104 d** | mut 48 | 0.92 | 619-624 | VQLIRN | AAAARN |
| **#104 e** | mut 49 | 0.83 | 619-624 | VQLIRN | AALIAA |
| **#104 f** | mut 50 | 0.96 | 619-624 | VQLIRN | VQAAAA |
| **#104 b** | mut 51 | 0.88 | 619-624 | VQLIRN | VQAARN |
| **#104 AQAA** | mut 52 | 0.94 | 619-624 | VQLIRN | AQAARN |
| **#104 VAAA** | mut 53 | 0.96 | 619-624 | VQLIRN | VAAARN |
| **#104 AAAN** | mut 54 | 0.96 | 619-624 | VQLIRN | VQAAAN |
| **#104 AARA** | mut 55 | 0.87 | 619-624 | VQLIRN | VQAARA |
| **#112 d** | mut 56 | 0.83 | 667-672 | CVVKPI | AAAAPI |
| **#112 f** | mut 58 | 0.83 | 667-672 | CVVKPI | CVAAAA |
| **#119 d** | mut 59 | 0.77 | 709-714 | PFCSHH | AAAAHH |
| **#119 e** | mut 60 | 0.96 | 709-714 | PFCSHH | AACSAA |
| **#119 f** | mut 61 | 0.93 | 709-714 | PFCSHH | PFAAAA |
| **#119 c** | mut 62 | 0.97 | 709-714 | PFCSHH | PFCSAA |
| **#119/120** | mut 63 | 0.79 | 709-720 | PFCSHHFNKLHL | AAAAAAAAAAAA |
| **#120 d** | mut 64 | 0.90 | 715-720 | FNKLHL | AAAAHL |
| **#120 e** | mut 65 | 0.95 | 715-720 | FNKLHL | AAKLAA |
| **#120 f** | mut 66 | 0.89 | 715-720 | FNKLHL | FNKLAA |
| **#122 d** | mut 67 | 0.93 | 727-732 | VVPCRH | AAAARH |
| **#122 e** | mut 68 | 0.84 | 727-732 | VVPCRH | AAPCAA |
| **#122 f** | mut 69 | 0.92 | 727-732 | VVPCRH | VVAAAA |
| **#122 b** | mut 70 | 0.70 | 727-732 | VVPCRH | VVAARH |
| **#122 c** | mut 71 | 0.64 | 727-732 | VVPCRH | VVPCAA |
| **#140 d** | mut 72 | 0.87 | 835-840 | WTDIPY | AADIAA |
| **#140 e** | mut 73 | 0.89 | 835-840 | WTDIPY | WTAAAA |

Excluded mutations for next step
